# *In Silico* insight to identify Potential Inhibitors of BUB1B from Mushroom bioactive compounds to Prevent Breast cancer metastasis

**DOI:** 10.1101/2022.06.14.496027

**Authors:** Divya Mishra, Ashish Mishra, M.P. Singh

**Affiliations:** Centre of Bioinformatics, Institute of Interdisciplinary Studies, University of Allahabad, Prayagraj (211002), Uttar Pradesh, INDIA; Centre of Biotechnology, Institute of Interdisciplinary Studies, University of Allahabad, Prayagraj (211002), Uttar Pradesh, INDIA

**Keywords:** Lepiotaprocerin D, Molecular docking, Molecular dynamics simulation, BUB1B, ADMET, Aneuploidy, Tumorigenesis

## Abstract

BUB1B (BUB1 Mitotic Checkpoint Serine/Threonine Kinase B) gene belongs to the spindle assembly checkpoint protein family, proven to be associated with many kinds of cancers. Protein kinases are crucial in metaphase to anaphase cellular transition. It is a primary element of spindle assembly checkpoint (SAC). This SAC is responsible for guarding the proper chromosomal segregation while any chromosomal instability hinders its normal functioning. The chromosomal instability may lead to cancer progression. And it occurs mainly due to overexpression of BUB1B in many cancer types especially in breast cancer metastasis. This could be controlled by designing proper inhibitor of this gene. For this purpose, there are proved existence of various mushroom compounds possessing some important medicinal properties with proven anti-cancerous effects. Proper screening and identification of molecules having these anti-cancerous effects and showing affinity against BUB1B with inhibition properties could be obtained from these compounds. So this study has incorporated 70 bioactive compounds (handpicked through literature mining) of distinct mushrooms that were considered and explored for identifying a suitable drug candidate. Their ADME/T properties were obtained to predict the drug-likeness of these 70 mushroom compounds based on Lipinski’s rule of 5 (RO5). The screening of these bioactive compounds and subsequent molecular docking against BUB1B provided compounds with the best conformation-based binding affinity. The best 2 complexes, i.e., BUB1B- Lepiotaprocerin D and BUB1B-Peptidoglycan, were subjected to the molecular dynamics simulation. Both the complexes were observed for their affinity, stability, and flexibility in protein-ligand complex systems. The MD simulation study revealed that Lepiotaprocerin D has an energetically favorable binding affinity with BUB1B. Results showed that the formation of a hydrogen bond between ASN123, SER157 residue and Lepiotaprocerin D has strengthened the affinity of Lepiotaprocerin D with BUB1B. Hence, this study identified Lepiotaprocerin D as a potential and novel inhibitor for BUB1B that could be a plausible drug candidate for checking and controlling the spread of breast cancer metastasis.

## 1 Introduction

The spindle assembly checkpoint (SAC) is used by the cells during mitosis to prevent segregation defects from unattached or improperly coupled chromosomes (Wang et al. 2022). SACs remain active on the kinetochore until they are firmly attached to the apparatus of the spindle. Finally, SAC satisfaction results in the release of CDC20 from inhibitory mitotic regulators, enabling the activation of the anaphase-inducing complex/cyclosome (APC / C) (Lee et al. 2017). Once activation is done by CDC20, the APC/C E3 ubiquitin ligase targets the degradation of several proteins, including securin, which ultimately leads to sister chromosome separation, and triggers the onset of anaphase. The SAC activity of kinetochore is driven by a network of protein interactions and the activities of various protein kinases, including Aurora B, MPS1, and Bub1b. MPS1 phosphorylated MELT (Met-Glu-Leu-Thr core consensus) repeats of knl1 kinetochore protein permits SAC to be stimulated by the recruitment of other SAC proteins, such as Bub1b (Lee et al. 2017). Aurora B is present in centromere with a combination of Haspin-mediated H3 histone phosphorylation (H2pT3) and Bub1b-mediated phosphorylation at Thr120 (H2ApT120), where it is necessary to elevate appropriate kinetochore binding and manage the SAC. As described in Aurora B, Bub1b plays a dual role in SAC and chromosome alignment. Although its contribution to the alignment of a chromosome has been consistently identified, analyses have generated conflicting results related to the requirement of Bub1b for SAC (Bolanos-Garcia et al. 2011). RNAi knockdown from HeLa and RPE1 and generation of conditional knockout mouse embryonic fibroblasts found Bub1b to be significant for SAC. In contrast, the initial CRISPR-CAS9 genome modification methods in RPE1 and HAP1 cells indicated only a minor role for Bub1b in SAC when the cells were susceptible through the inhibition of Mps1. These contradictory results were primarily reconciled with the discovery of nonsense-associated alternative splicing that allows some Bub1b expression followed by CRISPR-CAS9 and that siRNA knockdown of Bub1b residues greatly impaired the SAC response in BUB1b disrupted cells. Recently, HAP1 cells with multiple Bub1b exons absent from genetic DNA were generated using two lead RNAs for CRISPER-CAS9. Surprisingly, SAC remained viable in these cells even when the more general approach was combined with the Bub1b siRNA knockdown. However, complete destruction of BUB1b could only occur in haploid HAP1 cells, but not in some other cell lines.In spite of the controversy, this experimental system allows for an active evaluation of BUB1b (Han et al. 2015). Bub1b has a Bub3 binding domain where Bub1b is centralized to the kinetochore. Bub1b’s central site acts as a platform for placement through Bub1b on the RZZ complex, placement of Mad1 / 2 and Cdc20 in kinetochore, and is required for Bub1b SAC functionality (Jiao et al. 2021). Many reports have found that Bub1b kinase activity is important for SAC activation; while others suggested that phosphorylation of Cdc20 via Bub1b could directly contribute to APC / C inhibition. Similarly, there are conflicting reports as to whether the activity of Bub1b kinase is necessary to chromosomal alignment. Bub1b H2ApT120 phosphorylation places Sgo1 / 2 and Aurora B, and other proteins from the chromosome assembly complex (CPC), in the centromeres and may integrate binding error correction with SAC signals (Robbyn et al. 2016). Indeed, H2ApT120 is necessary to maintain centromeric SAC activity and Aurora B in the absence of H3pT3. In the end, Bub1b is also auto-phosphorylated, both within and outside the kinase domain of the activation moiety, which may play a role in controlling the localization of Bub1b (Qiu et al. 2020). The active properties of Bub1b would benefit from a low molecular weight kinase inhibitor. In fact, drugs targeting Mps1 and Aurora B are powerful tools for breaking down kinase activity and mitotic division. Potent and specific inhibitors of Bub1b kinase are particularly important because the complete elimination of genetic deletions or siRNAs in human cells is difficult and only the remaining 4% of Bub1b is required for SAC activity. However, in the light of many conflicting reports on the effectiveness of Bub1b, it is important to validate the inhibitors used to evaluate their effectiveness correctly. The availability of potential drugs against BUB1B may help to suppress the potential for cancer and its metastases, therefore molecular screening is necessary to identify potential molecules that may bind Bub1b and inhibit its activity. It is always useful and beneficial to use natural compounds as medicines because they are much safer in terms of their pharmacokinetics and dynamics properties. Studies have shown that various mushrooms have significant anti-cancer properties **(Rajewska et al., 2004)**.

Over the past two decades, there have been numerous reports of the anti-tumor properties of these most incredible mushrooms that have attracted the attention of the scientific community around the world **(Ajith et al., 2007; Sliva et al., 2008)**. Due to cost-effectiveness, natural appearance and negligible side effects of these mushrooms extract make them more favorable for medicinal use. *Pleurotus ostreatus* is one of the most cultivated and widespread macro funguses, with various medicinal properties, including anti-tumor, antioxidative, hypocholesterolemic, antiatherogenic activities (Gu et al., 2006; Bobek et al., 1995; Sarangi et al., 2006; Bobek et al., 1999). In the earlier 2008, Jedinak et al demonstrated that *P. ostreatus* have inhibition property against human colon and cancer (Jedinak et al., 2008); later P. ostreatus extract has also been studied for its carcinogenic potential in other cancers and found to be effective on erythroleukemia and human breast and gastric cancer (Tong et al., 2009; Ebrahimi et al., 2017; Facchini et al., 2014; Cao et al., 2015). Since the last decade, several structure based insilico approaches such as molecular docking, homology modeling and MD simulation, are already exist as potential tools for screening compounds against receptor having substantial accuracy (Katara, 2013; Kesharwani et al., 2021). As 3-D structure of BUB1B (PDB: 2WVI) is available (fig. 1), its create opportunities to perform structure based analysis for its molecular interaction. The current study aims to screen biomolecules from distinct variety of mushrooms to find out potential lead compounds against BUB1B target (Ma et al. 2017). For this purpose 70 already reported bioactive compounds of mushrooms having anticancer properties were utilized. All 70 compounds were screened for their pharmacokinetics (ADMET) properties, suitable compounds were screened against BUB1B through structure-based molecular interaction approach, i.e., molecular docking, and further, their stability and affinity were observed through molecular dynamics (MD) simulation.

**Figure 1.**
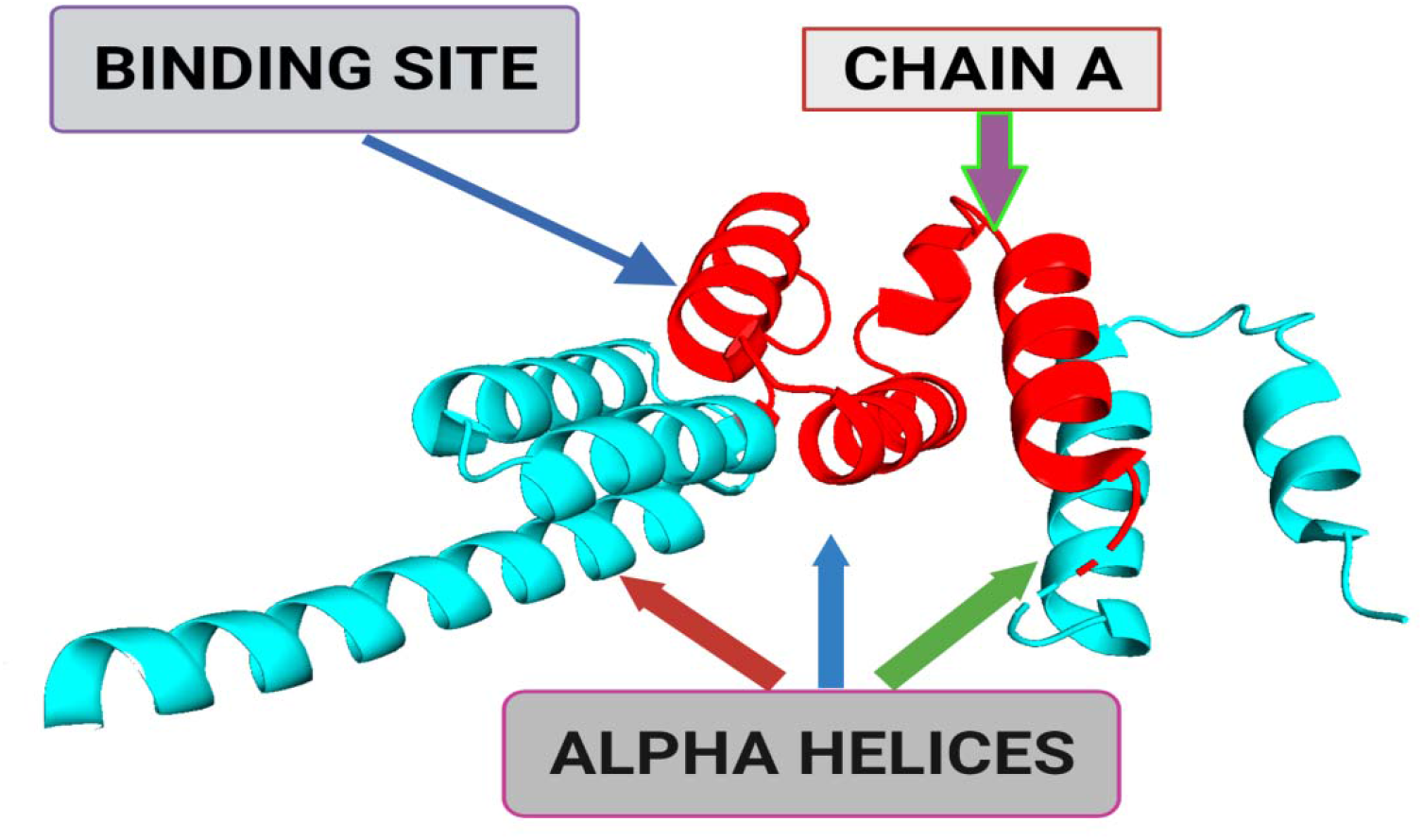
3-Dimensional structural representation of BUB1B (PDB ID: 2WVI; Resolution = 1.80 Å)

## 2 Materials and Methods

### 2.1 Screening and selection of Mushroom bioactive compound

The 70 bioactive compounds from edible mushrooms were obtained from different literature mining. The essentially predicted physicochemical descriptors of the selected compounds were obtained by calculating ADMET properties from SwissADME and helped in screening the compounds with favorable pharmacokinetic properties. A comprehensive analysis of the log P (octanol/water), and QP% (human oral absorption) were obtained from this SwissADME program. Another very important part which is essential for rational drug design is Lipinski’s rule of 5 (RO5) of the bioactive compounds and this was also obtained from SwissADME and also evaluated the acceptability of the known compounds.

### 2.2 Molecular interaction analysis through docking

For screening of selected bioactive compounds against BUB1B (PDB: 2WVI), AutoDock Vina (Trott et al., 2010) was employed. Amongst these bioactive compounds with favorable pharmacokinetic properties, top five compounds having best binding affinity scores were further filtered out and these were then docked against the BUB1B receptor. For obtaining the active site coordinates and residues of the receptor PyMol (v 2.5) was utilized. This process was based on the pdb file of receptor-ligand complex. PyMol provided the coordinates (x_center = 0.4537, y_center= -24.6765, z_center = -12.2435, x_size= 60, y_size = 50, z_size = 70) to obtain the grid parameter file and these coordinates belong to chain-A of the protein; 72-165 residues. Further, in the next step, the grid map files were obtained by running AutoGrid. These grid map files were then used for docking using AutoDock, v4.2.6 suite (Morris et al., 2009). There were several standard parameters set for the docking purpose. These included hybrid GA-LS runs = 50, number of individuals in population = 300, rate of gene mutation = 0.02, maximum number of generations = 27000, maximum number of energy evaluations = 2500000, rate of crossover = 0.8. The algorithm that was employed for AutoDock run was Lamarckian Genetic Algorithm (LGA) and it provided RMSD (Root mean square deviation) values and binding energies. Out of the five compounds, now the top two compounds having best docking score were further chosen for validation through molecular dynamics simulation.

### 2.3 Molecular dynamics (MD) simulation

Desmond v3.6 Package (Bowers et al., 2006) was employed for validating the findings of molecular docking using molecular dynamics simulation. This was done for elucidating the effectiveness of these mushroom bioactive compounds for the activation of BUB1B. For this purpose, the preparation of the lead compounds viz. lepitaprocerin D and peptidoglycan (muramic acid) with BUB1B protein was done using the OPLS2005 force field. Further, for building the system, the pre-defined TIP3P water model was used and these were acting as water molecules. The orthorhombic periodic conditions set at 10 Å units were used for constructing these. Before MD simulations, the energy minimization of the system was done, and also the balancing of Na+/Cl- was obtained to neutralize the charges of the complexes electrically. The steepest descent method was used as a part of minimization protocol for the complexes. The heating process was carried out at 0-300 K having time steps of 0.001 ps and annealing steps of 2000. The system further normalized at 1000 steps in an equilibrium state with 0.001ps time steps. Last step was the final production step of the system that continued up to 100ns at 300K temperature and 1 Atm pressure at 0.001ps time steps. These were applied by employing Nose-Hoover method with NPT ensemble. The pictorial representation of the methodology employed for the present study is shown in Fig. 2.

**Figure 2.**
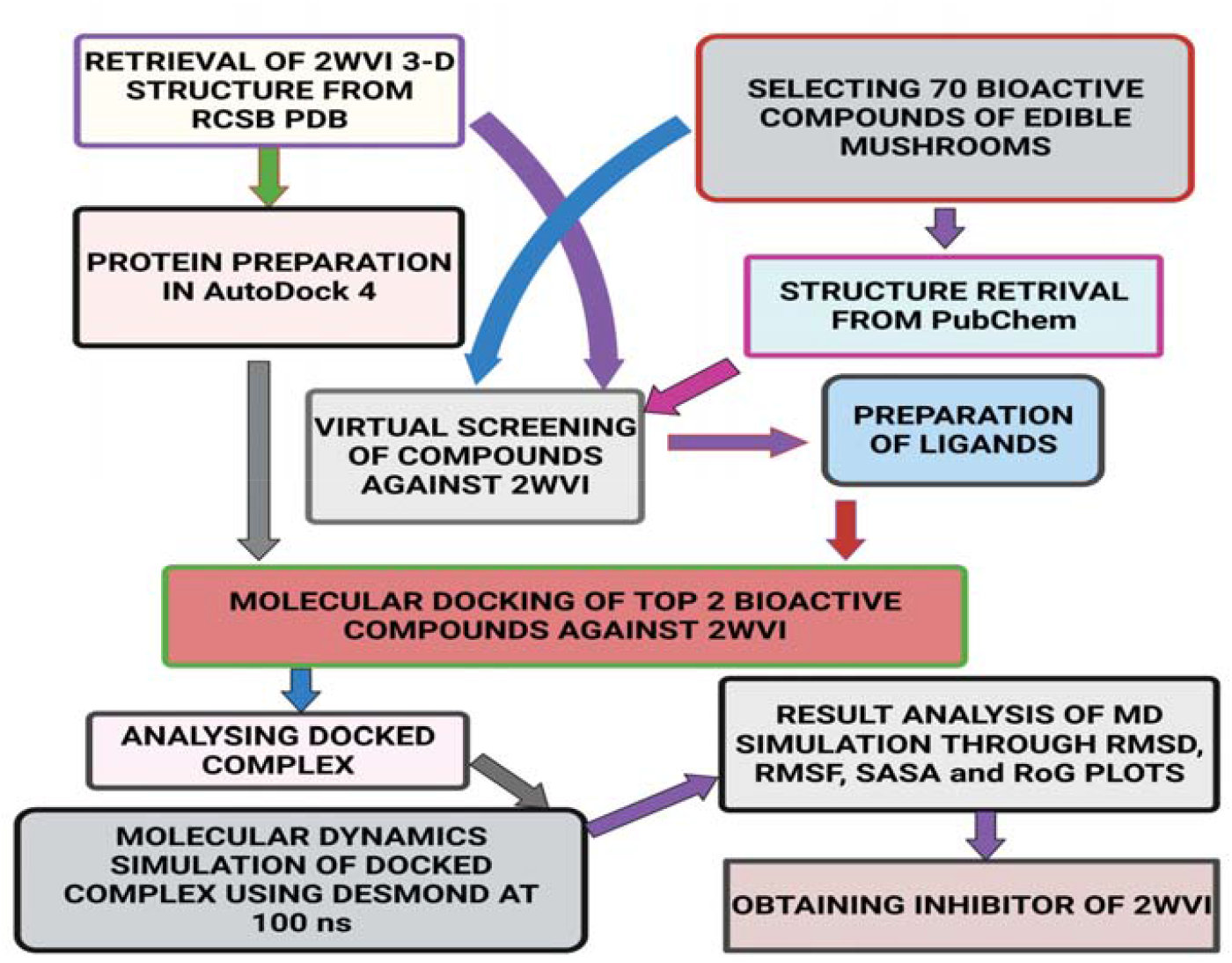
Schematic representation of the methodology followed in the present study.

## 3. Results

### 3.1 ADMET property Analysis

All compounds administrated through the oral route have to go through the absorption, distribution, metabolism, and excretion process. Before considering any bioactive compound as lead it is imperative to check its ADME/T features so that any compound with inappropriate pharmacokinetics properties must be filtered out (Guan et al., 2019). The drug-like activity of the selected HMW and LMW edible mushroom compounds was categorized using ADME/T properties shown in the table-1.

**Table 1.** ADME/T properties of selected 5 bioactive compounds.

| S. No. | Compound Name | Mol. Volume | Mol. Weight | Rotatable Bonds | VdW PSA | RO5 Voil | HB Donors | HB Acceptors |
| --- | --- | --- | --- | --- | --- | --- | --- | --- |
| 1 | Grifolin | 195.05 | 328.5 | 8 | 40.46 | 0 | 2 | 2 |
| 2 | Ganoderiol | 217.32 | 474.7 | 6 | 80.92 | 0 | 4 | 4 |
| 3 | Illudin | 121.00 | 264.3 | 1 | 77.76 | 0 | 3 | 4 |
| 4 | Lepiotaprocerin D | 210.37 | 480.6 | 2 | 72.83 | 0 | 1 | 5 |
| 5 | Peptidoglycan | 112.38 | 251.23 | 4 | 142.47 | 0 | 5 | 8 |

The physicochemical descriptors include molecular weight, molecular volume, H-bond donors, H-bond acceptors, and their position according to Lipinski’s rule of five. Lipinski’s rule of 5 is the rule of thumb to evaluate drug-likeness; if a chemical compound contains certain pharmacological and biological properties, then properties would make it an active drug. The rule describes molecular properties important for a drug’s pharmacokinetics in a living system, including ADMET properties. All these pharmacokinetic parameters are within the acceptable range defined for human use, thereby indicating their potential as drug-like molecules. Lepiotaprocerin D had two rotatable bonds with polar surface area (PSA) 72.83, whereas the number of rotatable bonds and PSA for peptidoglycan was 4 and 142.47, respectively.

### 3.2 Molecular interaction analysis through molecular docking

The receptor BUB1B (PDB: 2WVI) was docked with the 70 compounds obtained from the literature survey. The active site of the BUB1B (2WVI) receptor consisted of Thr85, Tyr81, Gln58, Asp79, Tyr89, Ser96, Ser157, Tyr139, and Glu161 amino acid residues. The screening with AutoDock Vina found five compounds, i.e., Grifolin, Ganodriol, Illudin S, Lepiotaprocerin D, and Peptidoglycan as most appropriate leads, as most appropriate leads **(Fig. 3)**.

**Figure 3:**
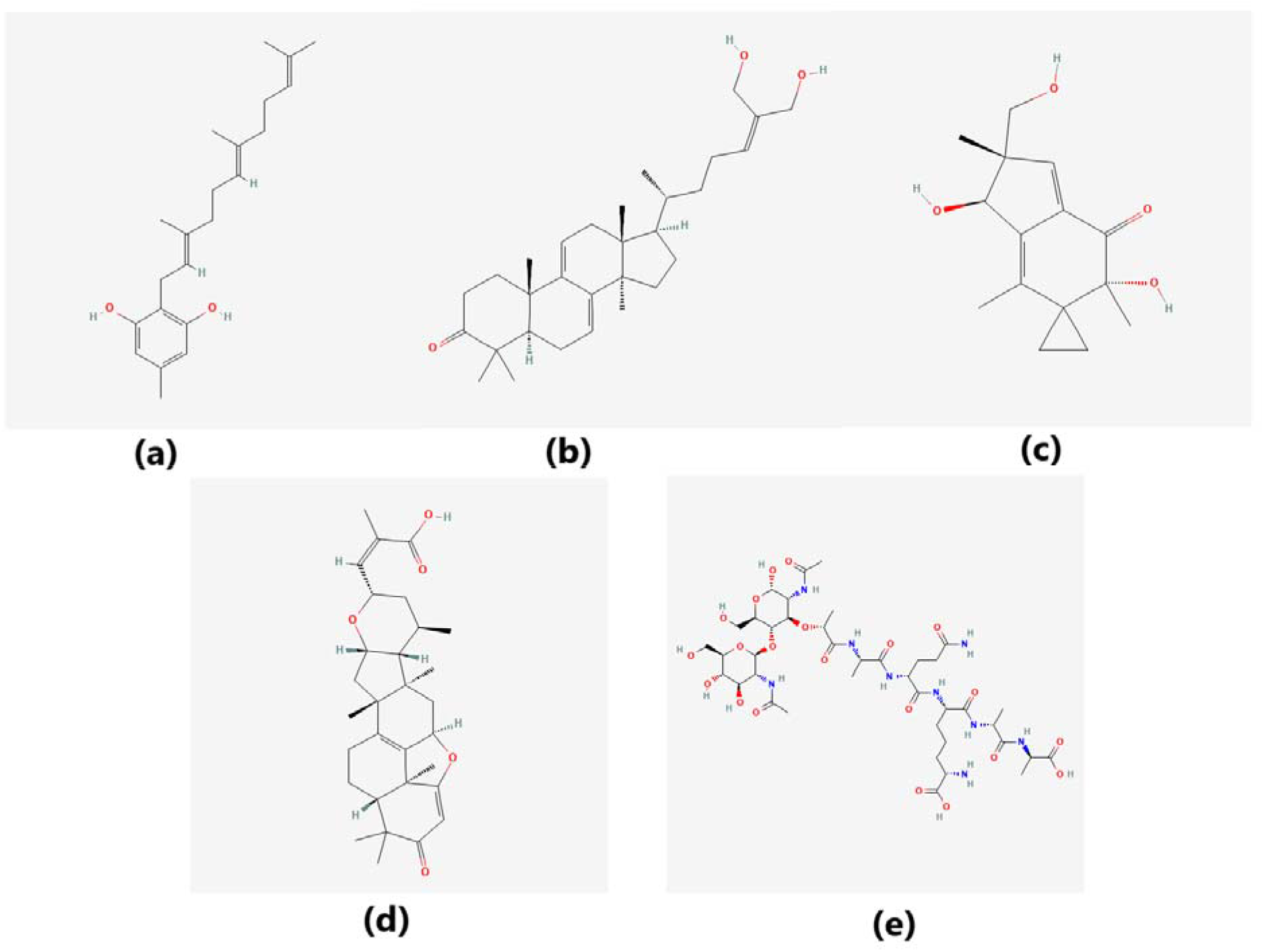
The five bioactive compounds that were docked with 2WVI receptor. (a) Grifolin (b) Ganderiol (c) Illudin S (d) Lepiotaprocerin D (e) Peptidoglycan.

These 5 compounds further docked against the BUB1B to obtain the best binding conformation and their affinity. The docking showed well-formed binding of the just identified lead compounds with one or more residues (amino acids) in the active site pocket of the 2WVI receptor shown in Fig. 3. This *in silico* molecular docking revealed that the newly identified compounds exhibit excellent binding energy towards the target receptor, as shown in the table (table 3).

**Table 3.** Top 5 bioactive compounds that have best binding affinity against BUB1B.

| S. No. | Compound | Binding Energy | RMSD | Partition Function | Free Energy (kcal/mol) | Internal Energy (kcal/mol) | Entropy (kcal/mol/K) |
| --- | --- | --- | --- | --- | --- | --- | --- |
| 1 | Lepiotaprocerin D | -8.91 | 37.33 | 50.70 | -2325.98 | -8.18 | 7.77 |
| 2 | Peptidoglycan | -8.70 | 37.58 | 50.63 | -2325.20 | -7.40 | 7.77 |
| 3 | Grifolin | -8.14 | 35.37 | 50.61 | -2325.00 | -7.20 | 7.77 |
| 4 | Ganderiol | -7.73 | 38.77 | 50.58 | -2324.62 | -6.82 | 7.77 |
| 5 | Illudin S | -7.08 | 35.19 | 50.60 | -2324.86 | -7.06 | 7.77 |

### 3.3. Molecular dynamic simulation analysis

Since docking only provides the static view of the interaction between the receptor and compound in the active site of the protein receptor, it doesn’t provide dynamic observation of interaction. Molecular dynamics simulation offers to compute of the binding atom movements with the time using newton’s motion. The molecular dynamics (MD) simulation was performed to find out the stability, confirmation, and intermolecular interaction of the top 2 ligand molecules (obtained from Autodock) with 2WVI protein as shown in Figure 4 (a) & (b). The time-dependent modification of the complexes was calculated over 100ns using the Desmond package. The MD simulation was performed under the thermodynamics conditions (applied volume, density, pressure, and temperature). The complete system was annealed and equilibrated using ensembles. Moreover, the final production step was performed to investigate the structural modification of the complex.

**Figure 4.**
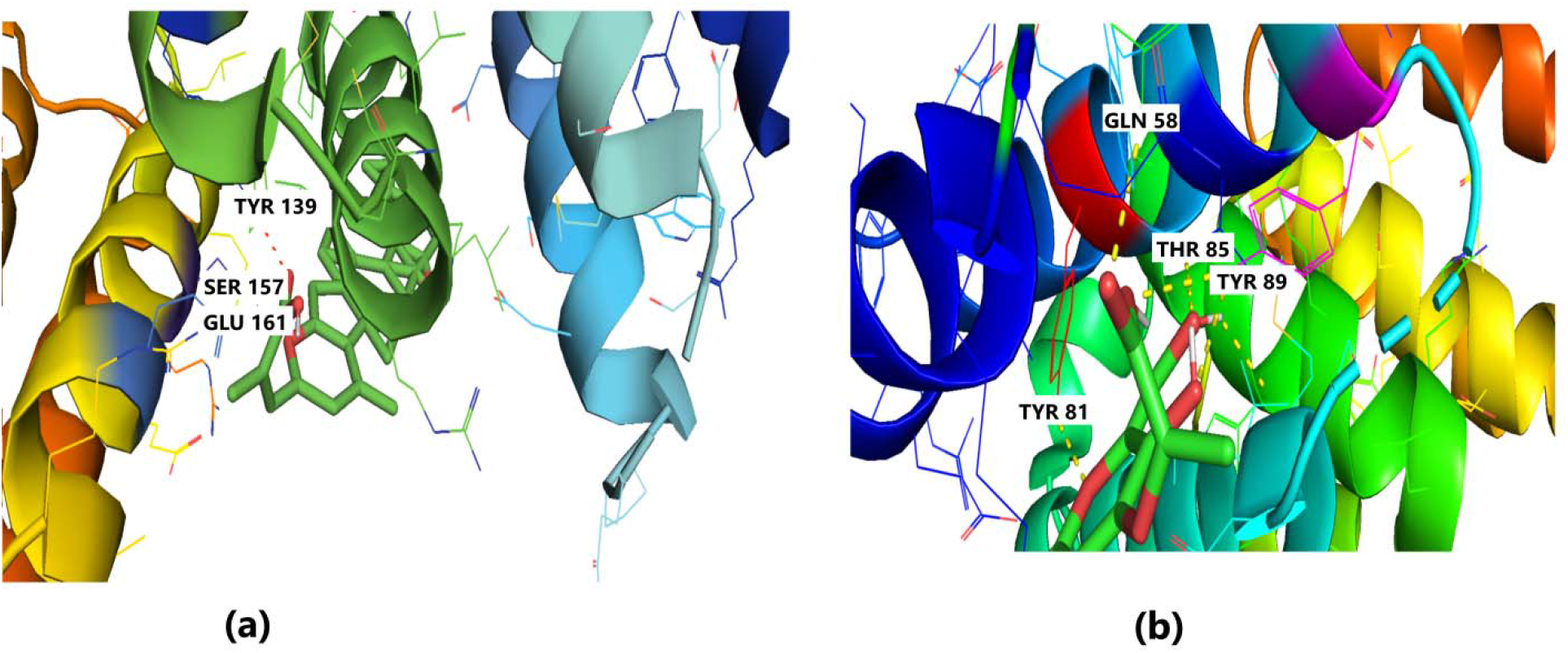
Molecular Docking of 2WVI with (a) Lepiotaprocerin D and (b) Peptidoglycan.

#### Root Mean Square Deviation (RMSD)

Root Mean Square Deviation (RMSD) is used to measure the deviation in the backbone of a protein from its initial conformation of structure to its final position. This deviation, which is produced during MD simulation, determines the protein’s stability relative to its conformation. Smaller deviation indicates a more stable protein structure. On the other hand, ligand RMSD indicates the stability of the ligand with respect to the receptor (protein) and its respective binding pocket. The RMSD value for the ligand should be significantly lower than the protein RMSD because if this value is large, it is possible that the ligand may diffuse away from its initial binding pocket. From Figure 5, it can be observed that for Lepiotaprocerin D, the system equilibrated after 40ns after which the fluctuations were stable till 6 Å. The ligand was stable with respect to protein and confined to the binding pocket of the protein till approximately 48ns after which the ligand diffused away from the binding pocket owing to its RMSD value becoming larger than that of the receptor but again returns back within the binding pocket after 52ns. Likewise, for peptidoglycan, the system gets equilibrated after 10ns after which fluctuations stabilize for a short span, and then again equilibration was obtained.

**Figure 5.**
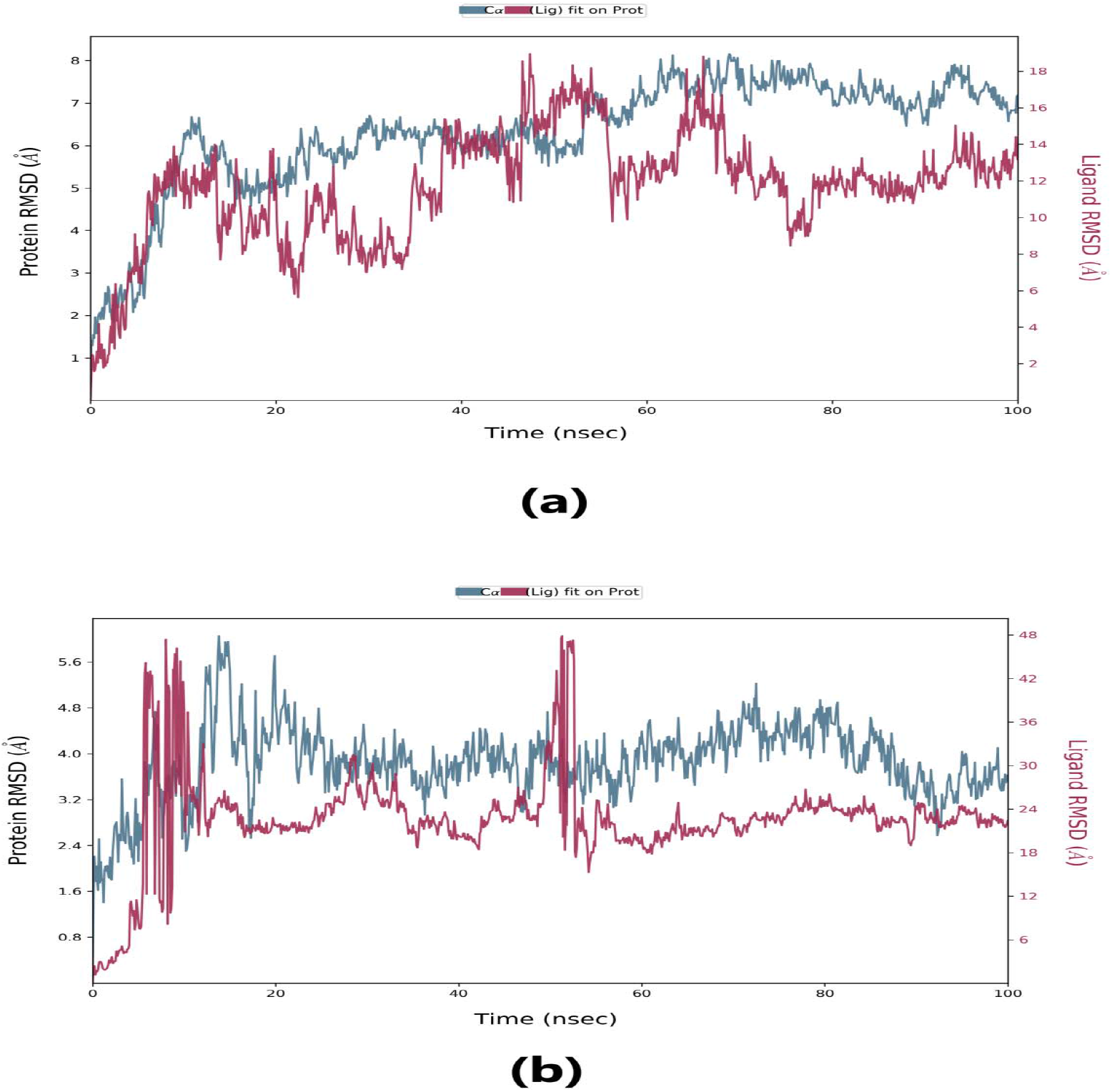
Root Mean Square Deviation (RMSD) plots of (a) Lepiotaprocerin D and (b) Peptidoglycan obtained from MD simulation in Desmond.

#### Root Mean Square Fluctuation (RMSF)

Root mean square fluctuation (RMSF) provides information about the regions that display higher flexibility levels. On the RMSF plot, peaks display the areas of maximum fluctuation of the protein. The Cα backbone for peptidoglycan displayed a higher peak at residue number 30, which indicates more movement and also showed the most flexible residues around residue number 120 to 140 Figure 6 (b). Similarly, in the case of lepiotaprocerin D, the highest peak for the Cα backbone, indicating more movement, was obtained at residue number 35. The most flexible residues were shown at residue numbers 40 to 80 shown in Figure 6 (a).

**Figure 6.**
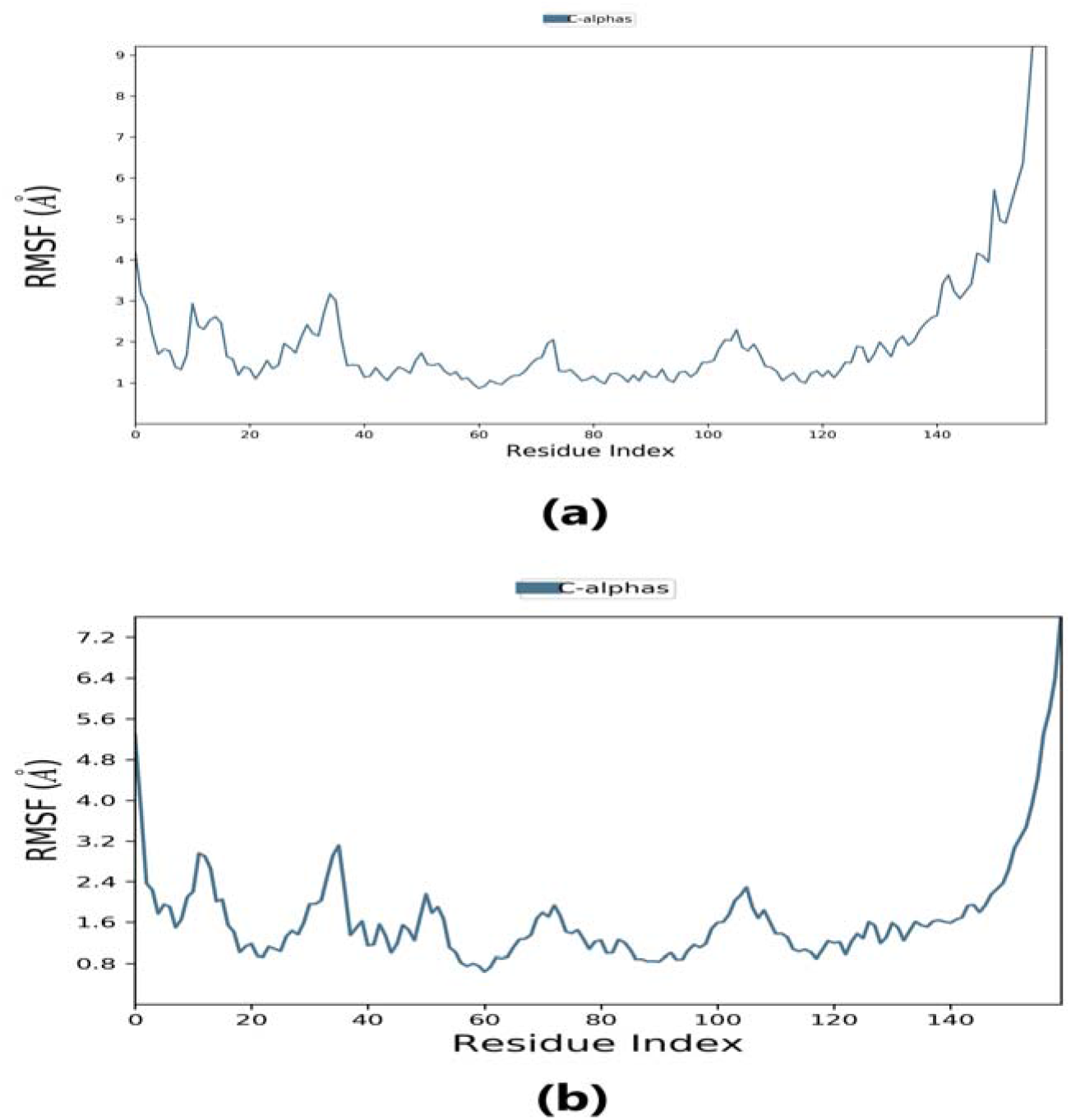
Root Mean Square Fluctuation (RMSF) plot of (a) Lepiotaprocerin D and (b) Peptidoglycan.

#### Protein-Ligand Contacts

Protein-ligand interaction was observed throughout the simulation. These contacts are basically of four kinds: hydrophobic, hydrogen bonds, ionic, and water bridges. These intermolecular interactions play a vital role at the atomic level for predicting the binding mode of lepiotaprocerin D and peptidoglycan with the 2WVI receptor. Lepiotaprocerin D forms a hydrogen bond with 2WVI along with binding site residues of Ser157 and Glu161, and water bridges along Tyr139, Ser157, and Glu161 shown in Figure 4 (a). Peptidoglycan, on the other hand, formed ionic interactions along Tyr81, hydrogen bond along Gln58, Tyr81, Thr85, and Tyr89, and water bridges along Gln58, Tyr81, and Tyr89. The hydrophobic interactions are relatively smaller in terms of interaction fraction as compared to those of lepiotaprocerin D. shown in Figure 4 (b). Also, lepiotaprocerin D contained three binding site residues where interactions occurred more than 10% viz. Tyr139, Ser157, and Glu161. However, there were no interactions of more than 10% in the case of peptidoglycan Figure 7 (a) & (b).

**Figure 7.**
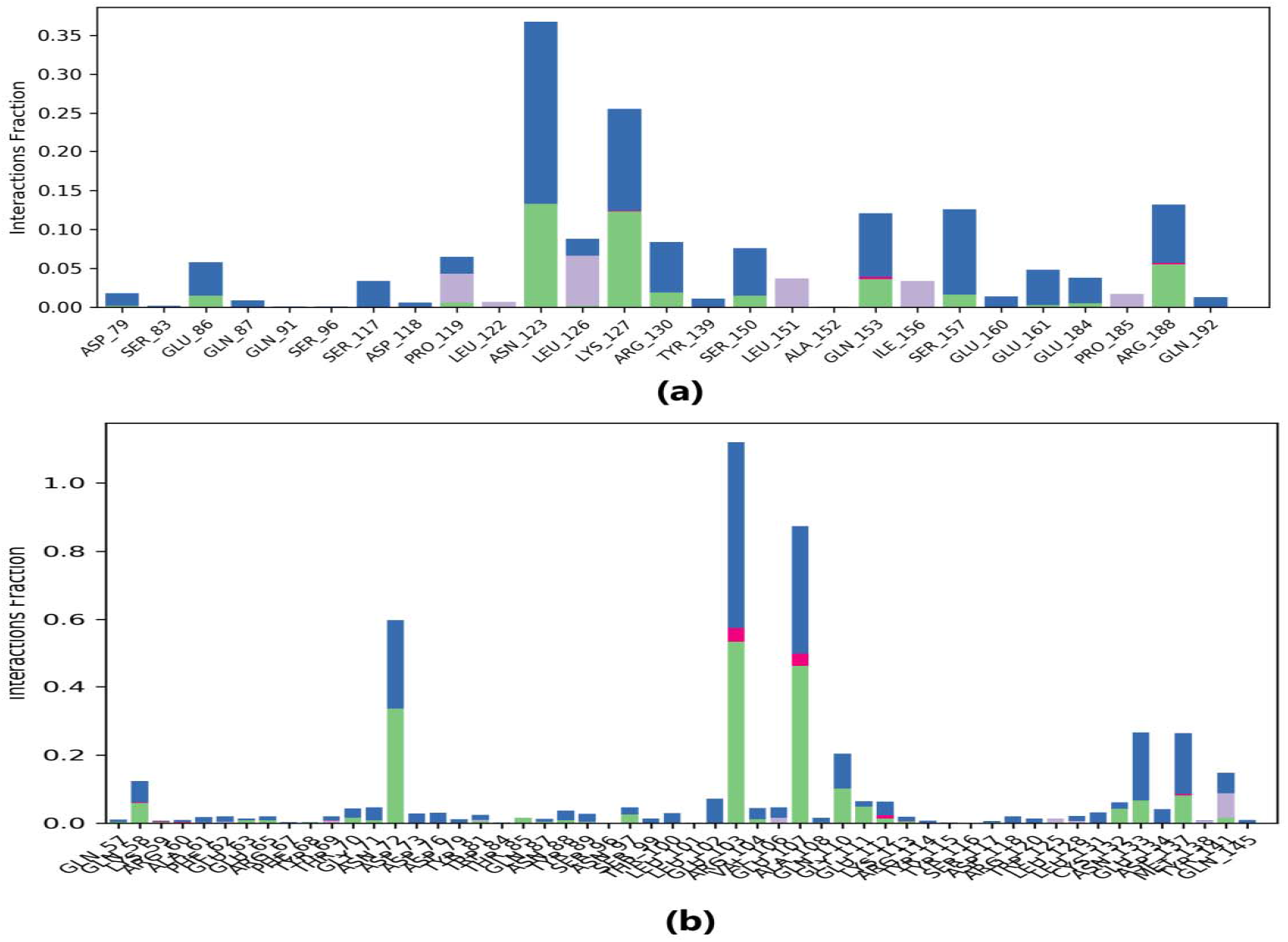
Protein-ligand contacts where blue color indicates water bridges, green color indicates H-bonds, grey color indicates hydrophobic bonds and pink color indicates ionic bonding. (a) PL contacts showing amino acid residues corresponding to interaction between 2WVI and lepiotaprocerin D (b) PL contacts showing amino acid residues corresponding to interaction between 2WVI and peptidoglycan.

#### Radius of Gyration (RoG) and Solvent Accessible Surface Area (SASA)

The solvent-accessible surface area analysis and radius of gyration analysis were carried out to ascertain the level of compactness in the structure, and the accessibility of the solvent of both lepiotaprocerin D and peptidoglycan. The radius of gyration of lepiotaprocerin D was observed from the figure as 19.8 Å (Figure 8 (a)) while that of peptidoglycan was 19.2 Å (Figure 8 (b)). SASA plot also showed a higher value for lepiotaprocerin D as compared to peptidoglycan shown in Figure 9 (a) & (b).

**Figure 8.**
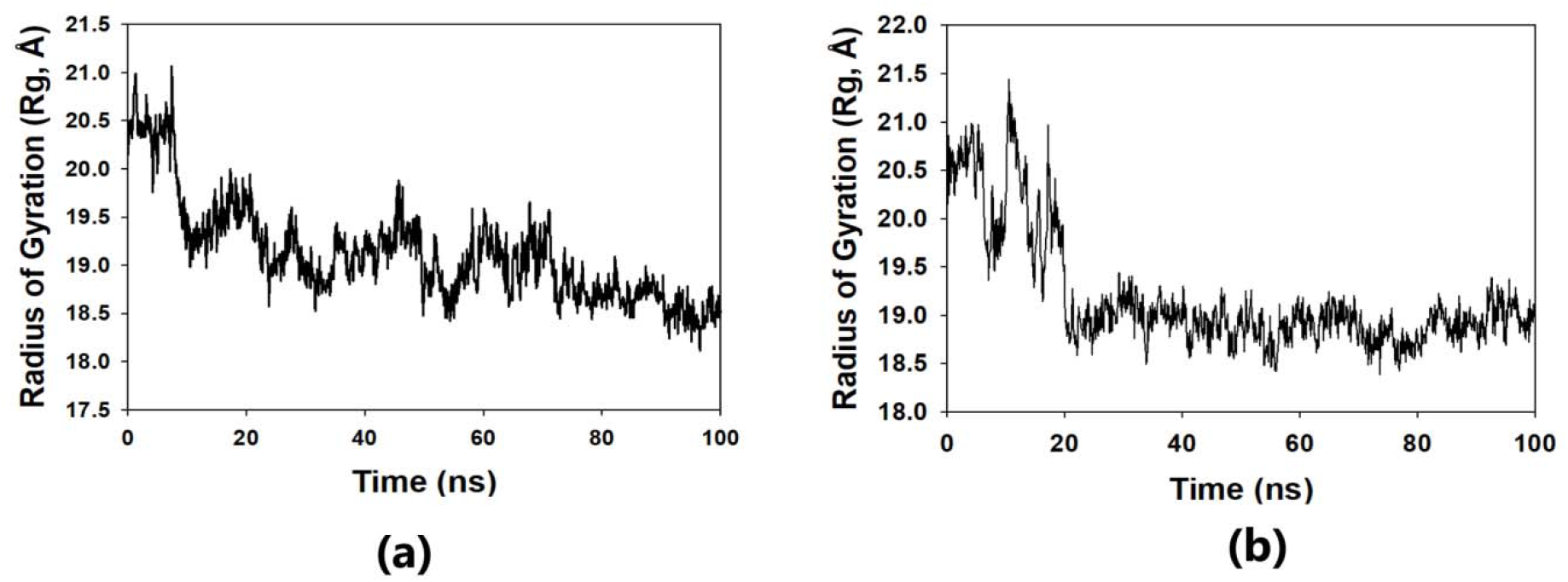
Radius of gyration plots corresponding to (a) Lepiotaprocerin D and (b) Peptidoglycan.

**Figure 9.**
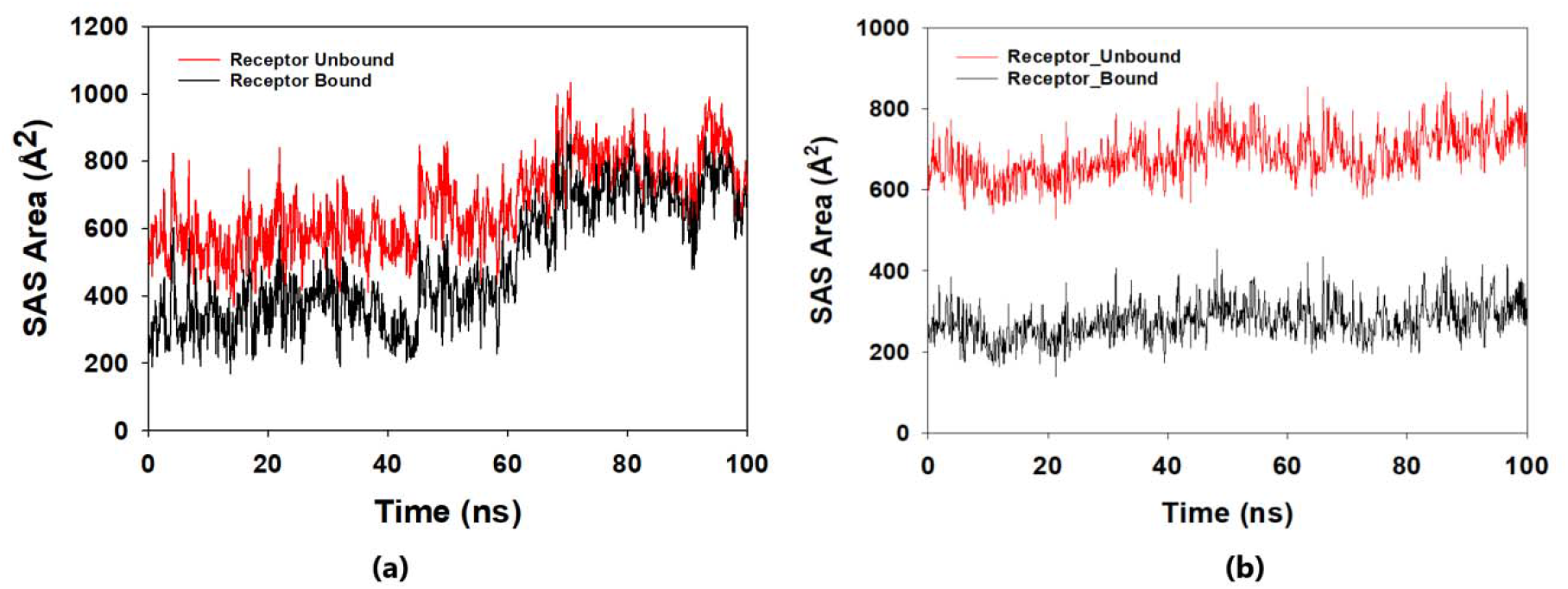
Solvent Accessible Surface Area (SASA) plots corresponding to (a) Lepiotaprocerin D and (b) Peptidoglycan.

## Discussion

The present study further focused on identifying the naturally occurring edible mushroom compounds like *P. ostreatus* compound that could be a potential anti-cancer drug and inhibit the BUB1B (2WVI) protein. The main cause of tumor progression is defective chromosomal segregation which is guarded by mitotic checkpoint (Holland et al., 2009). This chromosomal instability causes aneuploidy which plays a vital role in tumorigenesis and cancer metastasis (Daniel et al., 2011). The 2WVI protein, which is the primary component of the spindle assembly checkpoint (SAC), is responsible for corroborating proper chromosomal segregation. The over-expression of 2WVI in cancer cells as compared to normal cells regulates the process of tumorigenesis. Hence, 2WVI is considered a potential therapeutic target for the treatment of cancers as its inhibition could reduce the process of aneuploidy and tumorigenesis and metastasis. Extract obtained from edible mushrooms *P. ostreatus*, reportedly suppressed the cell proliferation of breast and colon cancers through p53-dependent as well as p53-independent pathways (Jedinak et al., 2008). For this study, the 70 bioactive compounds belong to different edible mushrooms extraction was taken, and their drug-likeness property was predicted based on their ADMET properties. All 70 bioactive compounds qualified Lipinski’s rule of 5 (RO5) and particularly Lepiotaprocerin D which has a molecular weight of 480.60 daltons (<500), HBD count of 1 (<5), and HBA count of 5 (<10). Hydrogen bond acceptor (HBA) and hydrogen bond donor (HBD) counts were considered important parameters for the permeability and polarity of compounds. Less value of HBD as against HBA indicated better efficacy and suitability of Lepiotaprocerin D as a potential drug. The molecular docking of 2WVI against these 70 compounds was carried out next to obtain the best conformations that could be reasonable for further studies. Through screening and docking, five bioactive compounds, i.e., Grifolin, Gandriol, Illudin S, Lepiotaprocerin D, and Peptidoglycan, were found to have the considerable binding affinity with 2WVI that make them potential inhibitors for overexpression of 2WVI. Two compounds, viz. Lepiotaprocerin D and peptidoglycan having the most negative affinities were further selected for molecular dynamics study, for 100ns simulation length, to obtain the dynamic behavior of atoms which provide information about structural changes, complex stability flexibility, and affinity at the atom and residue level. The RMSD analysis showed that lepiotaprocerin D was more stable as compared to peptidoglycan as it has lower RMSD values as compared to those of 2WVI till approximately 40ns. The RMSF analysis, which was done to measure fluctuation corresponding to each amino acid residue, indicated that lepiotaprocerin D has comparatively lower RMSF values of residues that mean less flexibility and more stability (Pandey et al., 2019). Furthermore, the binding free energy for peptidoglycan (2325.20 kcal/mol) was weaker than that of lepiotaprocerin D (2325.98 kcal/mol). This was consistent with experimental results and proved the efficacy of lepiotaprocerin D as a potential drug candidate.

In this study, lepiotaprocerin D was bound with 2WVI along SER157, and this binding proposed this compound most suitable for inhibitor design. A novel discovery of this study was the formation of the hydrogen bond of ASN123 with lepiotaprocerin D that further strengthened the binding of 2WVI and lepiotaprocerin D as a potential inhibitor. Due to the highly helical nature of 2WVI in the case of lepiotaprocerin D, the binding between 2WVI and lepiotaprocerin D was found to be more stable. The radius of gyration that evaluates the compactness of protein structure was shown to be decreasing in the case of lepiotaprocerin D, which gives an insight of more and more stable conformation of the protein 2WVI. The decrease in the solvent accessibility surface area (SASA) in the case of lepiotaprocerin D supported our observation for the radius of gyration. Hence, these parameters of molecular dynamic simulations supported the efficacy of lepiotaprocerin D for the treatment of different cancers and could be implied for the design of a suitable inhibitor that could reduce the effect of over-expressed 2WVI, thereby reducing tumorigenesis and metastasis of breast cancer and also the progression of glioblastoma multiforme (GBM) and hepatocellular carcinoma (HCC).

The limitations of this study are that the above discussed research work was carried out using *in silico* approach and hence the results can be validated through wet lab approach to obtain the significance of the obtained results.

## 5. Conclusion

BUB1B (BUB1 Mitotic Checkpoint Serine/Threonine Kinase B) is a primary element of the spindle assembly checkpoint. As overexpression of BUB1B is related to various cancers and its metastasis, thus there is a need for suitable bioactive compounds which can inhibit it without any side effects. For this purpose 70 bioactive compounds from edible mushroom which possesses some medicinal qualities and has proven anti-cancerous properties, have been screened against BUB1B. Structure-based *in-silico* approaches, i.e., virtual screening, docking, and MD simulation have been used. Two potential bioactive compounds lepiotaprocerin D and peptidoglycan were found to shows good drug-likeness and static affinity within binding cavity of BUB1B through docking. Further dynamics study, through MD simulation (100ns; RMSD, RMSF, RoG, and SASA), revealed that lepiotaprocerin D has a considerably good binding affinity with BUB1B and forms energetically favorable, stable, complex with BUB1B. Therefore, the study report lepiotaprocerin D as a potential and novel inhibitor for BUB1B that could be a plausible drug candidate. As lepiotaprocerin D belongs to edible mushroom, its use as an inhibitor drug against BUB1B can be safer and more effective.

## Acknowledgements

The authors are grateful to the Centre of Bioinformatics and Centre of Biotechnology, University of Allahabad for providing the access to the resources that were required to carry out this research work.

## Ethics approval and consent to participate

Not applicable

## Consent for publication

Not applicable

